# HCV infection activates the proteasome via PA28γ acetylation and heptamerization to facilitate the degradation of RNF2, a catalytic component of polycomb repressive complex 1

**DOI:** 10.1101/2024.06.02.596761

**Authors:** Hirotake Kasai, Atsuya Yamashita, Yasunori Akaike, Tomohisa Tanaka, Yoshiharu Matsuura, Kohji Moriishi

## Abstract

We previously reported that hepatitis C virus (HCV) infection or HCV core protein expression induces HOX gene expression by impairing histone H2A monoubiquitination via a proteasome-dependent reduction in the level of RNF2, a key catalytic component of polycomb repressive complex 1 (PRC1) (J. Virol, 2021, 95, e01784-20). In this study, we aimed to investigate the mechanism by which HCV infection accelerates RNF2 degradation. Yeast two-hybrid screening and an immunoprecipitation assay revealed that RNF2 is a PA28γ-binding protein. The proteasome activator PA28γ destabilized the RNF2 protein in a proteasome-dependent manner, since RNF2 degradation was impaired by PA28γ knockout or MG132 treatment. HCV infection or core protein expression reduced the levels of RNF2 and histone H2A K119 monoubiquitination and induced the expression of HOX genes in the presence of PA28γ, while PA28γ knockout reversed these changes. Treatment with a lysine acetyltransferase inhibitor inhibited the acetylation of PA28γ at K195 and the degradation of the RNF2 protein, while treatment with a lysine deacetylase inhibitor accelerated these events in a PA28γ-dependent manner. RNF2 protein degradation was increased by expression of the acetylation mimetic PA28γ mutant but not by expression of the acetylation-defective mutant or the proteasome activation-defective mutant. Furthermore, HCV infection or core protein expression facilitated the interaction between PA28γ and the lysine acetyltransferase CBP/p300 and then accelerated PA28γ acetylation and heptamerization to promote RNF2 degradation. These data suggest that HCV infection accelerates the acetylation-dependent heptamerization of PA28γ to increase the proteasomal targeting of RNF2.

**IMPORTANCE:** HCV is a causative agent of HCV-related liver diseases, including hepatic steatosis, cirrhosis and hepatocellular carcinoma. PA28γ, which, in heptameric form, activates the 20S core proteasome for the degradation of PA28γ-binding proteins, is responsible for HCV-related liver diseases. HCV core protein expression or HCV infection accelerates RNF2 degradation, leading to the induction of HOX gene expression via a decrease in the level of H2Aub on HOX gene promoters. However, the mechanism of RNF2 degradation in HCV-infected cells has not been clarified. The data presented in this study suggest that PA28γ acetylation and heptamerization are promoted by HCV infection or by core protein expression to activate the proteasome for the degradation of RNF2 and are responsible for HCV propagation. This study provides novel insights valuable for the development of therapies targeting both HCV propagation and HCV-related diseases.

## INTRODUCTION

Hepatitis C virus (HCV) is a major human pathogen that is currently causing chronic infection in 50 million people worldwide, with approximately 1.0 million new infections occurring annually, and puts infected individuals at risk of hepatic steatosis, liver cirrhosis, and hepatocellular carcinoma (HCC) (1). The development of direct-acting antiviral agents (DAAs) against HCV has led to significant improvements in the treatment of HCV infections over the past decade (2, 3). However, it has also been reported that hepatitis C patients remain at risk of HCC even after viral elimination (2). Understanding the pathogenesis of HCV infection remains an important challenge for the development of effective hepatitis C therapies. HCV belongs to the genus *Hepacivirus* in the *Flaviviridae* family and possesses an enveloped nucleocapsid containing a positive-sense RNA genome with a length of 9.6 kb. The single polyprotein (∼ 3000 amino acids) encoded by the viral genome is cleaved by viral and host proteases into 10 final viral proteins, which are classified as structural or nonstructural proteins (4). The structural proteins are the core, E1 and E2 proteins, and the nonstructural proteins are the p7, NS2, NS3, NS4A, NS4B, NS5A, and NS5B proteins. Nonstructural proteins play important roles in viral genome replication, assembly, and budding, whereas structural proteins, together with the host lipid bilayer, form viral particles (4). The core protein, a component of the nucleocapsid, has been implicated in the formation of viral particles as well as the induction of HCV-related pathological changes (4). Mice expressing the core protein in the liver were found to exhibit HCV-related liver diseases such as type II diabetes mellitus (5), hepatic steatosis (6, 7), and HCC (8, 9), suggesting that the core protein plays an important role in inducing HCV-related pathological changes.

We previously reported that the HCV core protein can promote a reduction in the monoubiquitination of lysine (K) 119 of histone H2A in the promoter regions of HOX alleles and subsequently activate the transcription of HOX genes (10). HOX family genes, a set of 39 genes divided into 4 clusters, namely, HOXA, HOXB, HOXC and HOXD, regulate body plan development during the neonatal stage and also regulate the growth, survival, migration and invasion of cells to support cancer development (11, 12). Monoubiquitination of H2A at K119 is catalysed by polycomb repressive complex 1 (PRC1) to repress target genes (13). The expression of the HCV core protein was found to increase the degradation of the RNF2 protein, a main catalytic component of PRC1, and reduce the level of K119-monoubiquitinated histone H2A (H2Aub) on the promoter regions of HOX alleles (10). The core protein-dependent degradation of RNF2 was found to be independent of polyubiquitination and to be inhibited by treatment with the proteasome inhibitor MG132 (10). However, the molecular mechanism by which the core protein induces polyubiquitination-independent, proteasome-dependent degradation of RNF2 remains unclear.

The ubiquitin‒proteasome system is a major proteolytic pathway that regulates various intracellular functions. The proteasome complex consists of the 20S proteolytic core and several proteasome activators (PAs), including the 19S PA (PA700), 11S PA (PA28 or REG), and PA200. The 19S PA can bind to the 20S proteasome and activate polyubiquitination-dependent proteolysis (14), while the 11S PA can bind to the 20S proteasome and activate polyubiquitin-independent proteolysis (14). The 11S PA is classified as a PA28α/PA28β heteroheptamer or a PA28γ homoheptamer. The expression of PA28α and PA28β is induced by interferon (IFN)-γ for the processing of MHC class I peptides (14). The PA28α/PA28β heteroheptamer activates the chymotrypsin-like activity of the 20S core proteasome (CP) in the cytoplasm, while the PA28γ homoheptamer activates the trypsin-like activity of the 20S CP in the nucleus (15). Several proteins, such as AID, SRC-3, p19, the HCV core protein and SirT1, have been reported to be substrates of the PA28γ-dependent proteasome system (4, 16–18). The amino acid sequence of human PA28γ is identical to that of its murine homologue and shares 55% identity with that of its tick homologue (19), suggesting that PA28γ is highly conserved from insects to mammals. This broad conservation indicates that PA28γ may regulate fundamental functions across the animal kingdom, although its detailed biological function is largely unknown. We previously reported that PA28γ binds to the HCV core protein and promotes the degradation of unfolded or nonfunctional core proteins through a polyubiquitin-independent proteosomal pathway (20, 21). Furthermore, PA28γ knockout abolished core protein-related liver defects (type II diabetes, steatosis, and HCC) in mice, suggesting that PA28γ plays a critical role in core protein-induced pathological changes (22, 23). Phenotypic analysis of PA28γ-knockout mice revealed that they were viable and capable of mating but exhibited mild growth defects (24), decreased sperm fertility (25), and accelerated senescence (26), although a critical function of PA28γ has not yet been identified.

In this study, we identified RNF2 as a novel PA28γ-binding protein and examined the effect of PA28γ on the stability of RNF2, monoubiquitination of histone H2A, regulation of HOX gene transcription, and propagation of HCV. We also aimed to clarify the mechanism by which the HCV core protein regulates PA28γ-dependent proteasome activity to inhibit PRC1 activity in HCV-infected cells.

## RESULTS

### PA28γ interacted with RNF1 and RNF2, components of polycomb repressor complex 1

To identify novel PA28γ-binding molecules, we screened a human foetal brain library using a yeast two-hybrid system with PA28γ as bait. We identified four positive clones, Cl-26, Cl-49, Cl-50 and Cl-53. Cl-26 included the gene encoding RNF1 in the open reading frame of GAL4-DNA binding domain, while Cl-49 included the gene encoding PA28γ in the frame. Cl-50 and Cl-53 included partial regions of the gene encoding pyruvate dehydrogenase complex component X isoform, but they did not match the frame. RNF1, a component of PRC1, was identified as a PA28γ-binding partner by this yeast two-hybrid screen. Since RNF2 is a catalytic component of PRC1 and is a homologue of RNF1, the RNF2 gene was isolated from total cDNA of the human liver. Plasmids encoding N-terminal FLAG-tagged PA28γ (FLAG-PA28γ), N-terminal HA-tagged RNF1 (HA-RNF1) and N-terminal HA-tagged RNF2 (HA-RNF2) were constructed to examine the interactions of PA28γ with RNF1 and RNF2 by immunoprecipitation in 293T cells. HA-RNF1 or HA-RNF2 was coimmunoprecipitated with FLAG-PA28γ, while FLAG-PA28γ was reciprocally coimmunoprecipitated with HA-RNF1 or HA-RNF2 (Fig. 1A). Next, we examined the subcellular localization of PA28γ and RNF2 by immunofluorescence staining. Unfortunately, we could not find an antibody that reacts specifically with endogenous RNF1. However, PA28γ and RNF2 were partially colocalized in the nucleus (Fig. 1B). The fluorescence intensities of PA28γ (green) and RNF2 (red) within each pixel were quantified and plotted (Fig. 1C), revealing a significant negative correlation for the colocalization of these two proteins (R = -0.856, Fig. 1C). These data suggest that PA28γ binds to RNF2 but is not strongly colocalized with RNF2 in untreated cells.

**FIG 1:**
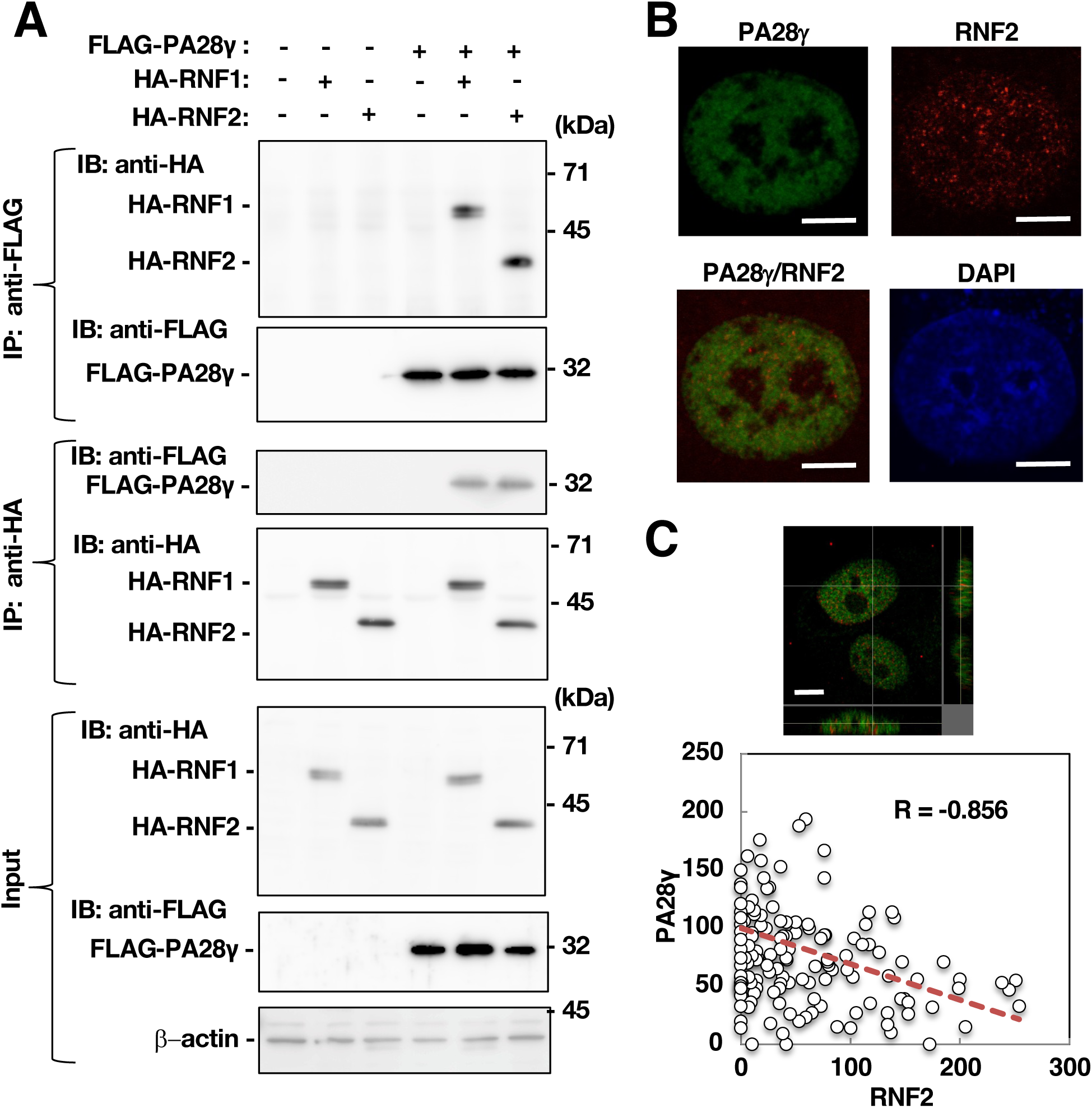
PA28γ could bind RNF2 but was not colocalized with RNF2. (A) FLAG-PA28γ was expressed in 293T cells together with HA-RNF1 or HA-RNF2. The cells were harvested 48 h post-transfection. These cell lysates were subjected to immunoprecipitation. Immunoprecipitates recovered with anti-HA or anti-FLAG antibodies were analysed by Western blotting using anti-FLAG or anti-HA antibodies, respectively. (B) The intracellular localization of PA28γ and RNF2 was examined by immunocytofluorescence. Images were acquired with a Keyence fluorescence microscope. White bars indicate 5 µm scale. (C) The image in the field of view shown in the top of panel C was acquired with a fluorescence microscope and subjected to three-dimensional reconstruction. The green and red fluorescence intensities in each pixel (in coordinate form) were calculated using the analysis software. Pearson’s correlation test was performed, and the correlation coefficient (R) was calculated. The data shown in this figure are representative of three independent experiments.

### PA28γ reduced the stability of RNF1 and RNF2

We hypothesized that the intracellular segregation of PA28γ and RNF2 may be due to PA28γ-dependent degradation of RNF2 following the interaction between these two proteins, because PA28γ-binding proteins are reportedly degraded via a polyubiquitin-independent proteosomal pathway (15). Using the CRISPR-Cas9 system, we established a PA28γ-knockout cell line and then examined the stability of the RNF1 and RNF2 proteins via a cycloheximide (CHX) chase assay. PA28γ was detected in wild-type Huh7OK1 cells, which are highly permissive to HCV infection (27), but not in PA28γ-knockout Huh7OK1 cells (Fig. 2A, B). The level of RNF2 in the wild-type cells was reduced by less than 50% 2 h after CHX treatment, whereas no significant reduction in the RNF2 level was observed in the PA28γ-knockout cells (Fig. 2A, B). RNF1 tended to be less stable in PA28γ-knockout cells than in wild-type cells, but the difference was not significant (Fig. 2A, B). In addition, the stability of BMI1 (PCGF4), another component of PRC1, was not affected by PA28γ knockout (Fig. 2A, B). These data suggest that PA28γ destabilizes RNF2.

**FIG 2.**
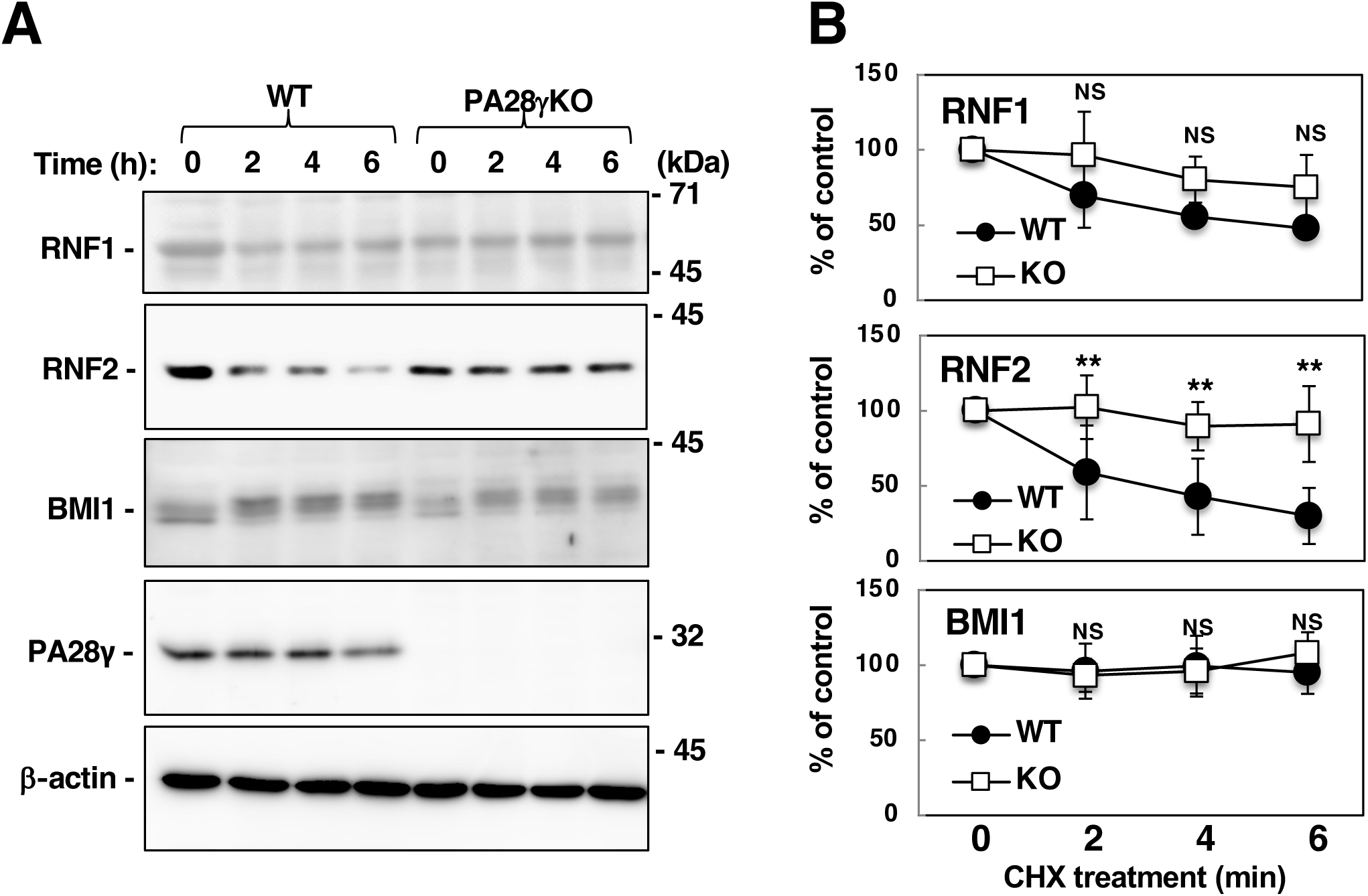
The endogenous RNF2 protein was stabilized by genetic knockout of PA28γ. (A) PA28γ gene knockout was conducted using the Huh7OK1 cell line and the CRISPR-Cas9 system. PA28γ was detected in the parental cells but not in the PA28γ-knockout cells. PA28γ-knockout Huh7OK1 cells or parental Huh7OK1 cells (Huh7OK1) were treated with the translation inhibitor CHX (2 mg/ml). Endogenous RNF1, RNF2 and BMI1 expression was evaluated by Western blotting. (B) The band densities of RNF1, RNF2, BMI1 and β-actin shown in panel A were estimated using ImageJ software. The values for RNF1, RNF2 and BMI1 were normalized to that of β-actin and are presented as percentages with respect to those in mock-infected cells. The p-value of each time measurement of KO cell line compared to that of WT cell line were lower than 0.01. The p-values of 2, 4, and 6 h measurements compared to 0 h measurement indicated 0.07, 0.02, and 0.01, respectively (the middle graph), suggesting that RNF2 was decreased in WT cells in a time-dependent manner. The data shown in this figure are representative of three independent experiments and are presented as the means ± SDs (n = 3). **: *p* < 0.01.

When PA28γ expression was restored in PA28γ-knockout cells, the endogenous RNF1 and RNF2 protein levels decreased in a PA28γ dose-dependent manner (Fig. 3A). However, the RNF1 and RNF2 mRNA levels were not affected by restoration of PA28γ expression (Fig. 3B). Furthermore, restoring the expression of PA28γ reduced the levels of HA-RNF1 and HA-RNF2 (Fig. 3C). HA-RNF1 and HA-RNF2 was expressed separately in 293T cells and were then isolated by immunoprecipitation using an anti-HA antibody for *in vitro* proteasome assays. The immunoprecipitates containing HA-RNF1 or HA-RNF2 were mixed with recombinant 20S CP with or without the 11S PA consisting of PA28γ. The levels of HA-RNF1 and HA-RNF2 were reduced in the presence but not in the absence of the 11S PA. These results suggest that PA28γ is involved in the proteolysis of RNF1 and RNF2. Since the RNF2 protein was more strongly destabilized by PA28γ than was the RNF1 protein, we subsequently focused on the relationship between RNF2 and PA28γ.

**FIG 3.**
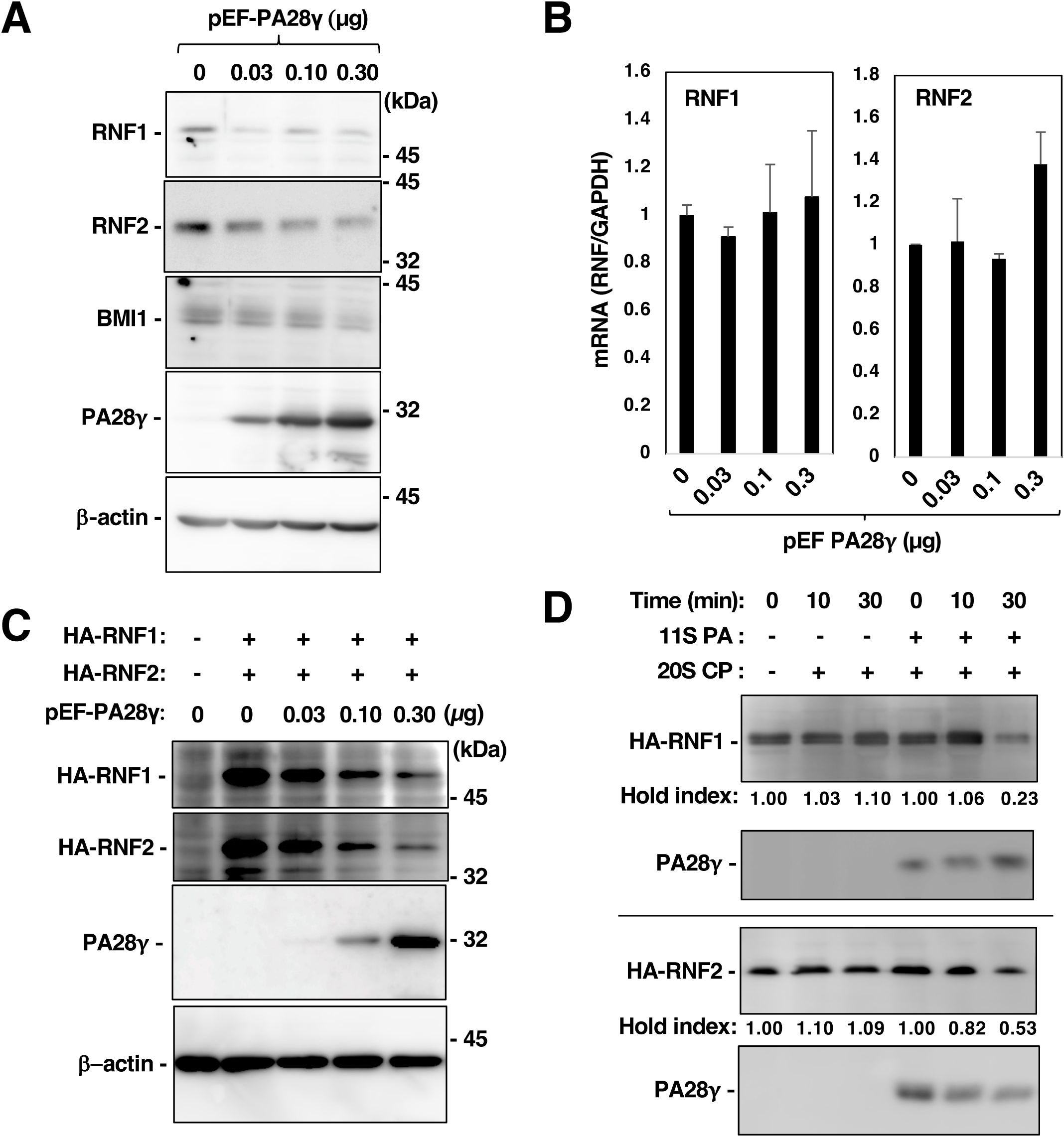
RNF2 degradation was promoted by the expression of PA28γ. (A) PA28γ-knockout Huh7OK1 cells were transfected with pEF PA28γ (0, 0.03, 0.1 and 0.3 µg/sample) or the empty vector. The total amount of transfected DNA in pEF PA28γ-transfected cells was supplemented with the empty plasmid to a final amount of 0.3 µg per sample. Endogenous RNF1, RNF2, BMI1, PA28γ and β-actin expression was evaluated by Western blotting. (B) Total RNA was prepared from the harvested cells described in panel A. The levels of RNF1 mRNA and RNF2 mRNA were estimated via RT‒qPCR. These mRNA levels were normalized to the GAPDH mRNA level and are presented as relative values compared to the empty plasmid control. The data shown in this figure are representative of three independent experiments and are presented as the means ± SDs (n =3). (C) PA28γ-knockout Huh7OK1 cells were transfected with pEF PA28γ (0, 0.03, 0.1 and 0.3 µg/sample) or the empty vector together with the plasmid encoding HA-RNF1 or HA-RNF2 (0.2 µg/sample). The total amount of transfected DNA in pEF PA28γ-transfected cells was supplemented with the empty plasmid to a final amount of 0.5 µg per sample. (D) The immuno-precipitates including HA-RNF1 or HA-RNF2 were mixed with 20S CP and 11S PA (PA28γ) in the reaction buffer and then incubated at 0, 10 or 30 min. The resulting preparations were subjected into Western blotting. The data shown in this figure are representative of three independent experiments.

### PA28γ promoted proteasome-dependent degradation of the RNF2 protein

PA28γ increases the trypsin-like activity of the ubiquitin-independent proteasome (15). We next examined the effect of proteasome activity on RNF2 stability in the presence of PA28γ. The RNF2 protein level was reduced in PA28γ-knockout cells by restoration of PA28γ expression, while MG132 treatment suppressed the PA28γ-dependent decrease in the RNF2 protein level (Fig. 4A). The intracellular localization of PA28γ and RNF2 was evaluated in the presence and absence of MG132. RNF2 was partially colocalized with PA28γ in the absence of MG132, while it strongly colocalized with PA28γ in the presence of MG132 (Fig. 4B). A significant negative correlation for the colocalization of wild-type PA28γ and RNF2 was found in the cells (Fig. 4C, a left half). The PA28γ mutant in which proline 245 (Pro245) was replaced with tyrosine (Tyr; PA28γP245Y) could not activate the proteasome (21, 28). This mutation lost the negative correlation between PA28γ and RNF2 localizations (Fig. 4C, a right half). These data suggest that PA28γ promotes the proteolytic degradation of RNF2 in a proteasome-dependent manner.

**FIG 4.**
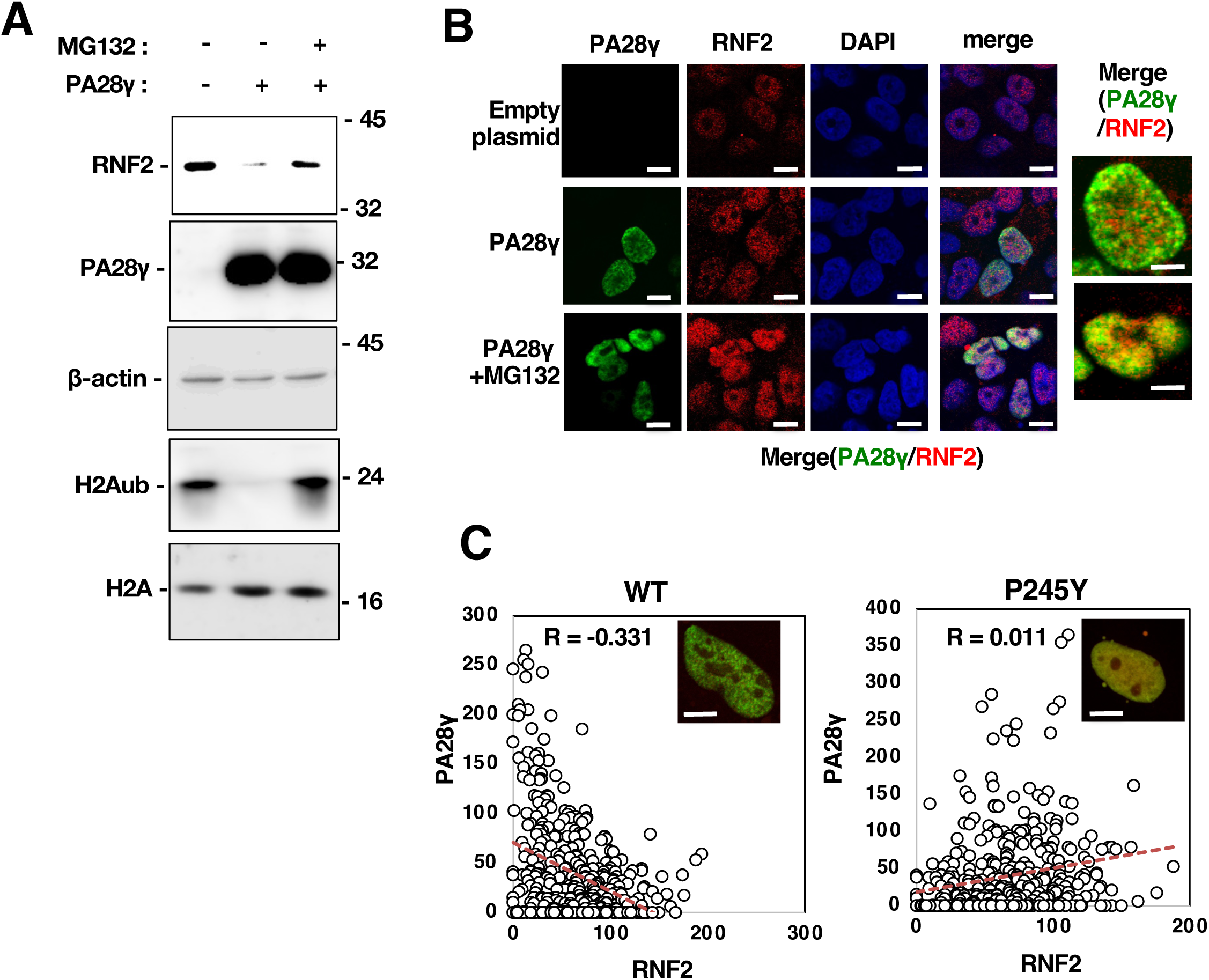
The degradation of the RNF2 protein and the reduction in H2Aub were promoted by PA28γ and suppressed by proteasome inhibition. (A) PA28γ-knockout Huh7OK1 cells were transfected with pEF PA28γ and treated with 10 µM MG132 at 48 h post-transfection. The resulting cells were harvested at 52 h post-transfection. These cell lysates were subjected to Western blotting. (B) The cells described in panel A were fixed and stained with the appropriate antibodies as described in the Materials and Methods. The intracellular localization of RNF2 and PA28γ was observed via confocal laser scanning microscopy. White bars in high magnified images (2 panels on right side) indicate 5 µm scale, while those in other panels indicated 10 µm scale. (C) PA28γ-knockout cells were transfected with plasmids encoding PA28γ or PA28γP245Y. The resulting cells were fixed 48 h post-transfection and stained with appropriate antibodies as described in the Materials and Methods. The nuclear localization of RNF2 and PA28γ was observed via Keyence fluorescence microscope. White bars indicate 5 µm scale. The green and red fluorescence intensities in each pixel (in coordinate form) were calculated using the analysis software. Spearman’s rank correlation test was performed. Each correlation coefficient (R) was shown. The data shown in this figure are representative of three independent experiments.

### PA28γ impaired the monoubiquitination of histone H2A at K119 and induced the transcription of HOX genes

PRC1 monoubiquitinates K119 of histone H2A to suppress the transcription of target genes via chromatin compaction (13, 29). RNF2 is an E3 ubiquitin ligase responsible for the catalytic activity of PRC1 for the monoubiquitination of histone H2A at K119 (13, 30, 31). We next examined the effects of PA28γ expression on histone H2A monoubiquitination and on the transcription of HOX genes. Restoration of PA28γ expression reduced the level of H2Aub (Fig. 5A) in PA28γ-knockout cells but did not affect the level of total histone H2A. We previously reported that HOXB9, HOXC13, and HOXD13 were completely silenced in Huh7OK1 cells but HOXA1 and HOXA3 was not (10). We evaluated the effect of PA28γ expression on the expression of the group with HOX gene silencing. Restoring the expression of PA28γ in PA28γ-knockout cells induced the transcriptional activation of HOXB9 (Fig. 5B). The level of H2Aub in the promoter region of HOXB9 was then evaluated by a chromatin immunoprecipitation (ChIP) assay and was found to be reduced by the restoration of PA28γ expression in PA28γ-knockout cells (Fig. 5C). The HOXB1, HOXC13 and HOXD13 genes were also transcriptionally activated by PA28γ expression (Fig. 5D), whereas HOXA1 and HOXA3 gene transcription was not affected by the restoration of PA28γ expression in PA28γ-knockout cells (Fig. 5D). These data suggest that PA28γ reactivates the expression of silenced HOX genes via impairment of H2A monoubiquitination.

**FIG 5.**
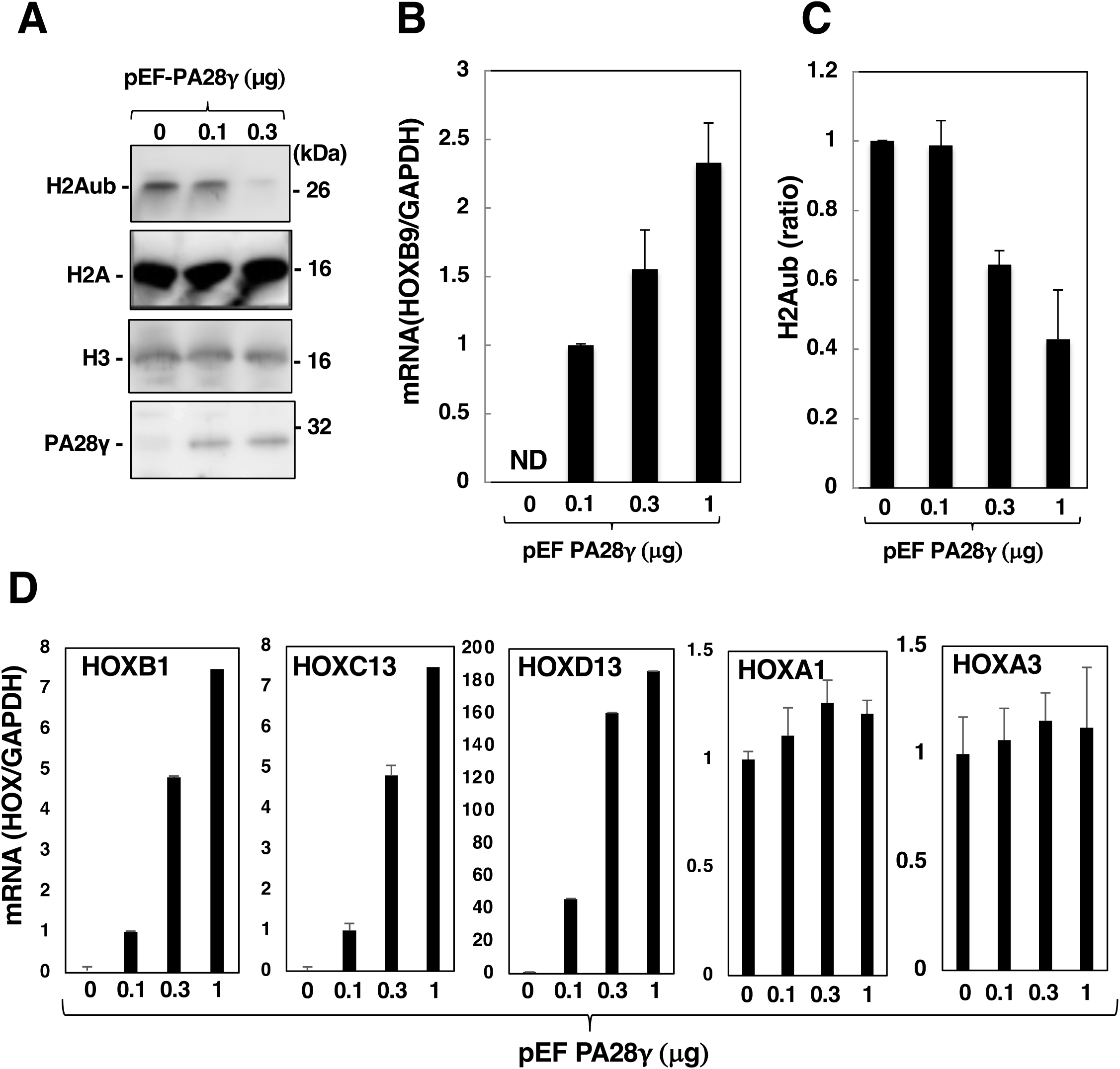
PA28γ was required for the induction of HOX genes. (A) PA28γ-knockout Huh7OK1 cells were transfected with pEF PA28γ (0, 0.1, 0.3 and 1.0 µg/sample) or the empty vector. The total amount of transfected DNA in these cells was supplemented with the empty plasmid to a final amount of 1.0 µg per sample. These cells were treated with 10 µM MG132 at 48 h post-transfection and were then harvested at 52 h post-transfection. These cell lysates were subjected to Western blotting. (B and D) Total RNA was prepared from the harvested cells described in panel A. The level of each HOX mRNA was estimated via RT‒qPCR. These mRNA levels were normalized to the GAPDH mRNA level and are presented as relative values compared to the empty plasmid control. The data shown in this figure are representative of three independent experiments and are presented as the means ± SDs (n =3). (C) The level of H2Aub in the promoter region of HOXB9 was evaluated by a chromatin immunoprecipitation (ChIP) assay. The data shown in this figure are representative of three independent experiments and are presented as the means ± SDs (n =3).

### HCV infection impaired the monoubiquitination of histone H2A at K119 and reduced the level of RNF2 in a PA28γ-dependent manner

We previously showed that HCV infection promoted RNF2 protein degradation via a polyubiquitination-independent proteasome pathway and subsequently reduced the level of H2Aub in the promoter regions of HOX genes (10). When wild-type Huh7OK1 cells were infected with HCV, the levels of H2Aub and RNF2 were reduced in a manner dependent on the number of days after infection in the presence but not in the absence of PA28γ (Fig. 6A, B). Knocking out PA28γ did not affect HCV replication, although it increased the level of the HCV core protein in cells and reduced the levels of both HCV RNA and infectious viral particles in the culture supernatant (Fig. 6C), consistent with the data shown in our previous report (21). These findings suggest that HCV infection impairs the monoubiquitination of histone H2A and the stability of RNF2 in a PA28γ-dependent manner.

**FIG 6.**
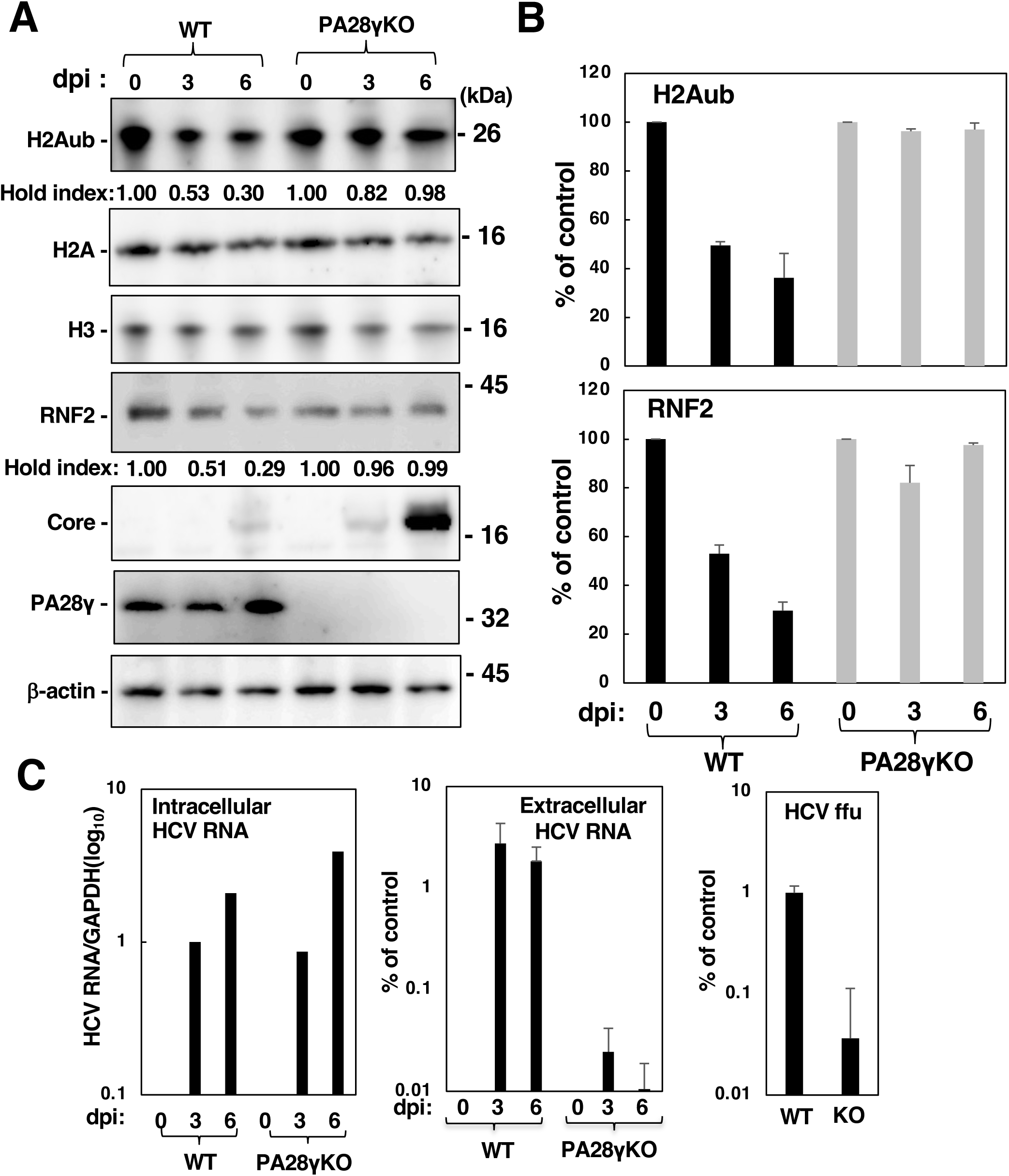
HCV infection reduced the levels of H2Aub and RNF2 in the presence but not in the absence of PA28γ. Huh7OK1 cells (WT) and PA28γ-knockout Huh7OK1 cells (PA28γKO) were infected with HCVcc at m.o.i. of 0.5 and harvested at 0, 3 and 6 dpi. These cell lysates were subjected to Western blotting (A), and the band densities of H2Aub and RNF2 were estimated by a method similar to that described in Figs. 2 and 5 (B). The levels of cellular and supernatant HCV RNA were estimated by RT‒qPCR, and the infectivity in the supernatant was estimated by a focus formation assay (C). The data shown in this figure are representative of three independent experiments and are presented as the means ± SDs (n =3).

### The HCV core protein reduced the level of H2Aub in the promoter regions of HOX genes and promoted their transcription in the presence but not in the absence of PA28γ

Our previous report suggested that polyubiquitination-independent proteolysis of RNF2 is promoted by the expression of the HCV core protein to activate HOX gene transcription in HCV-infected cells (10). Next, we examined whether the expression of the HCV core protein affects the RNF2 protein level, the H2A monoubiquitination level and HOX gene transcription in the presence or absence of PA28γ. Expression of the core protein reduced the levels of H2Aub and RNF2 in the presence but not in the absence of PA28γ, although the level of the core protein was markedly higher in PA28γ-knockout cells than in wild-type cells (Fig. 7A). A ChIP assay was carried out to estimate the level of H2Aub in the HOXB9 promoter region. The level of H2Aub in the HOXB9 promoter region was reduced by expression of the core protein in the presence but not in the absence of PA28γ (Fig. 7B). On the other hand, core protein expression promoted the transcription of the HOXB9 gene in the presence but not in the absence of PA28γ (Fig. 7C). These data suggest that the core protein reduces the levels of RNF2 and H2Aub and increases the expression of HOX gene in the presence but not in the absence of PA28γ.

**FIG 7.**
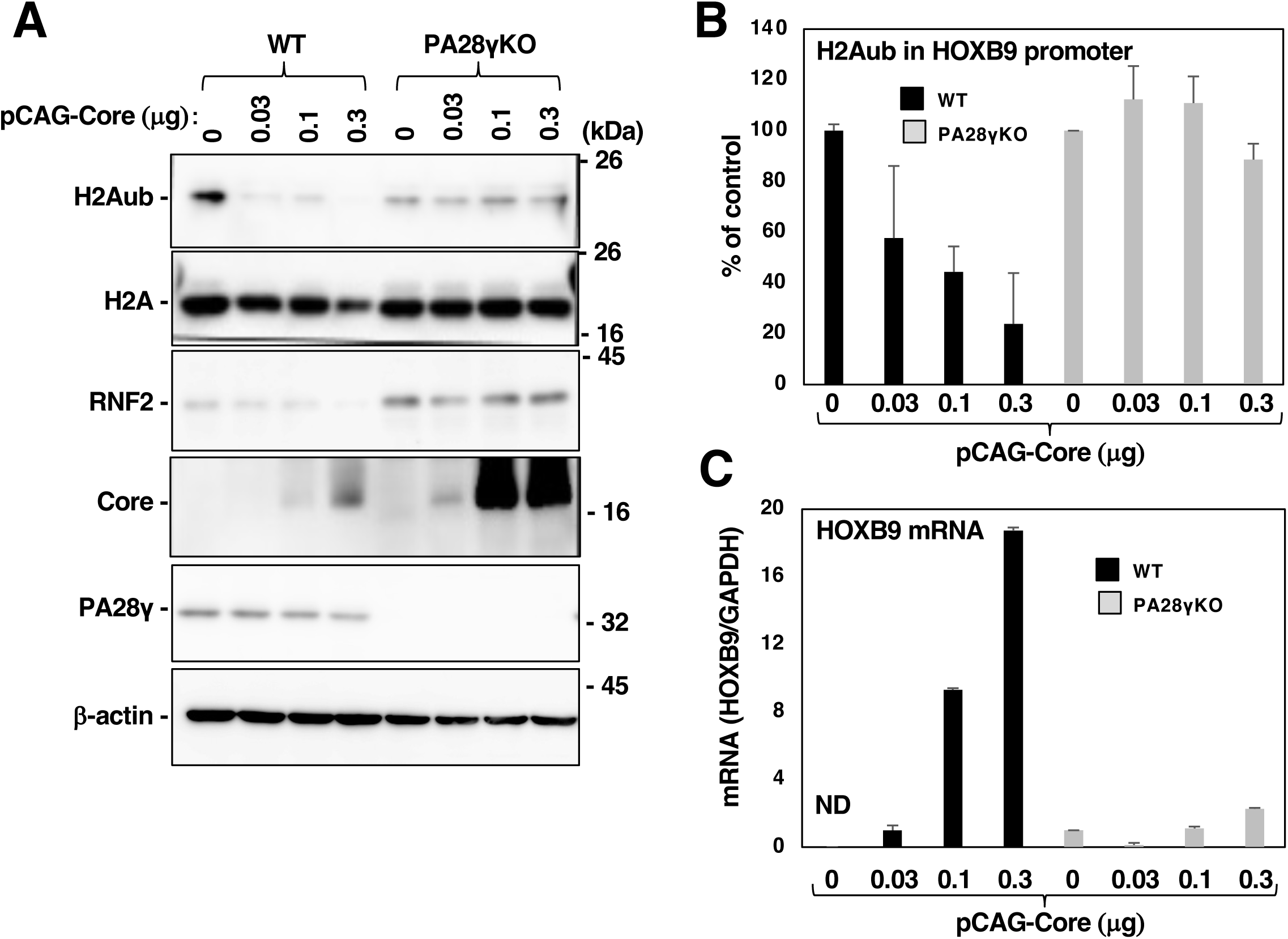
HCV core protein expression reduced the levels of H2Aub and RNF2 in the presence but not in the absence of PA28γ. (A) PA28γ-knockout Huh7OK1 cells (PA28γKO) and the parental cells (WT) were transfected with pCAG Core (0, 0.1, 0.3 and 1.0 µg/sample) or the empty vector. The total amount of transfected DNA in these cells was supplemented with the empty plasmid to a final amount of 1.0 µg per sample. These cell lysates were subjected to Western blotting. The data shown in this figure are representative of three independent experiments. (B) The cells described in panel A were subjected to a ChIP assay to estimate the level of H2Aub in the promoter region of HOXB9. The data shown in this figure are representative of three independent experiments and are presented as the means ± SDs (n =3). (C) Total RNA was prepared from the cells described in panel A. The level of HOXB9 mRNA was estimated by RT‒qPCR. The data shown in this figure are representative of three independent experiments and are presented as the means ± SDs (n =3).

### Acetylation of PA28γ at Lys195 promoted its homoheptamerization and RNF2 degradation

PA28γ was reported to be acetylated at Lys195 (K195) to promote its homoheptamerization for 20S CP binding and proteasome activation (32). CBP/p300, a lysine acetyltransferase (KAT), was reported to be responsible for the acetylation of PA28γ at K195 (32). Moreover, the core protein can interact with CBP and increase its acetyltransferase activity (33). We hypothesized that the core protein promotes CBP-dependent acetylation of PA28γ and augments the homoheptamerization of PA28γ to activate the proteasome to target RNF2 for degradation. We examined the effect of the KAT inhibitor C646 on PA28γ-dependent RNF2 degradation. Restoring the expression of PA28γ reduced the RNF2 level in PA28γ-knockout cells; in addition, CHX treatment promoted the reduction in the RNF2 level in the presence of PA28γ, whereas C646 treatment impaired the reduction in the RNF2 level in the presence of PA28γ regardless of CHX treatment (Fig. 8A). Immunoprecipitation using an anti-acetylated lysine (Ace-K) antibody and the lysates used to acquire the data shown in Fig. 8A showed that C646 treatment suppressed PA28γ acetylation (Fig. 8B). On the other hand, treatment with the lysine deacetylase inhibitor trichostatin A (TSA) promoted the decrease in the RNF2 level in the presence but not in the absence of PA28γ (Fig. 8C), while TSA treatment increased PA28γ acetylation (Fig. 8D). PA28γK195R, in which Lys195 is replaced with Arg, is an acetylation-defective mutant, while PA28γK195Q, in which Lys195 is replaced with Gln, is an acetylation mimetic mutant (32). A CHX chase assay was carried out to investigate the stability of RNF2 in the presence of wild-type PA28γ or these mutants. Six hours after treatment with CHX, the RNF2 level was 21% of that in the control cells in the cells expressing wild-type PA28γ, 104% of that in the control cells in the cells expressing PA28γK195R and 14% of that in the control cells in the cells expressing PA28γK195Q. These results suggest that the regulation of PA28γ acetylation controls the PA28γ-dependent degradation of RNF2. The HCV core protein increased the degradation of RNF2 in the absence but not in the presence of C646, suggesting that the core protein promotes the PA28γ-dependent degradation of RNF2 via KAT activation. These data suggest that HCV infection or the core protein expression promotes PA28γ acetylation for destabilization of RNF2.

**FIG 8:**
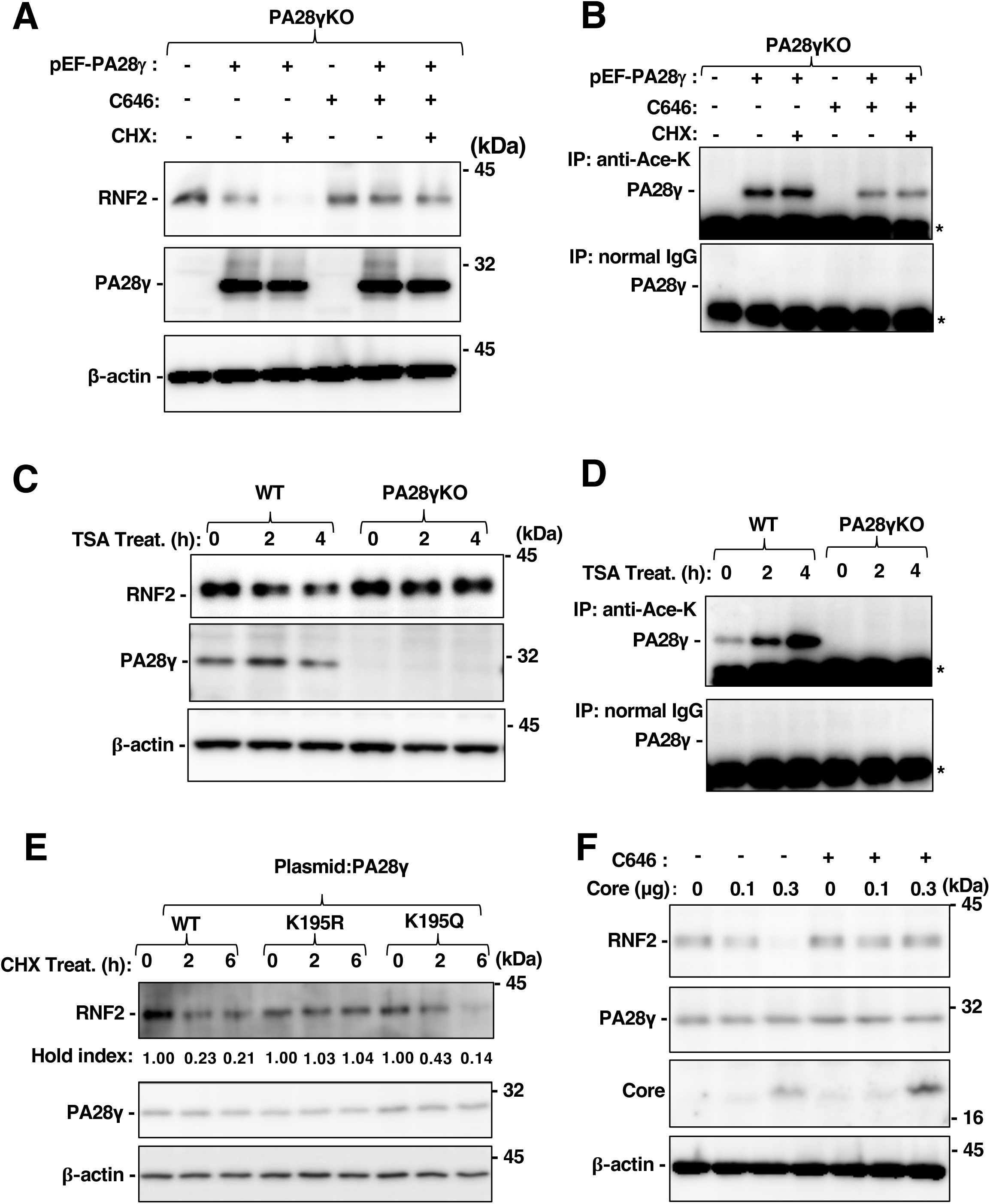
Acetylation of PA28γ at K195 was essential for the reduction in the RNF2 protein level. (A and B) PA28γ-knockout HuhOK1 cells were transfected with pEF PA28γ or the empty vector and were then treated with C646 and CHX at 36 h post-transfection. The cells were collected at 42 h post-transfection. These cell lysates were subjected to Western blotting. (C and D) PA28γ-knockout HuhOK1 cells were transfected with pEF PA28γ or the empty vector and were then treated with TSA at 36 h post-transfection. The cells were collected at 0, 2 and 4 h posttreatment. These cell lysates were subjected to Western blotting. (E) PA28γ, PA28γ K195R or PA28γK195Q were expressed in PA28γ-knockout Huh7OK1 cells. The transfected cells were treated with CHX and harvested at 0, 2 and 6 h posttreatment. These cell lysates were subjected to Western blotting. (F) Huh7OK1 cells were transfected with pCAG Core (0, 0.3 and 1.0 µg/sample) or the empty vector. The total amount of transfected DNA in these cells was supplemented with the empty plasmid to a final amount of 1.0 µg per sample. The cells were treated with CHX at 36 h post-transfection and harvested at 42 h post-transfection. These cell lysates were subjected to Western blotting.

### HCV infection promoted the acetylation of PA28γ via CBP activity

We investigated the mechanism by which HCV infection or core protein expression promotes PA28γ acetylation. Huh7OK1 cells were infected with HCV, and the infected cells were harvested at 0, 3 and 6 dpi. The cell lysates were subjected to immunoprecipitation using rabbit anti-Ace-K IgG or normal rabbit IgG. The levels of acetylated PA28γ and PA28γ homoheptamers increased in a time-dependent manner after infection (Fig. 9A, B). When PA28γ expression was restored with or without the expression of the core protein in PA28γ-knockout cells, the core protein promoted PA28γ acetylation and homoheptamerization (Fig. 9B, C), consistent with the data shown in Fig. 9A. The core protein has been reported to interact with CBP and subsequently increase its acetyltransferase activity (33). EGFP-PA28γ and FLAG-CBP were coexpressed in PA28γ-knockout cells, and EGFP-PA28γ coimmunoprecipitated with FLAG-CBP (lane 4, Fig. 9D). Furthermore, the core protein increased the amount of EGFP-PA28γ coimmunoprecipitated with CBP (lane 5, Fig. 9D). The RNF2 protein level was decreased in HCV-infected cells but was restored by CBP knockdown in infected cells (Fig. 9E). Taken together, these results suggest that the HCV core protein in infected cells reinforces the interaction between PA28γ and CBP, induces the acetylation and homoheptamerization of PA28γ, and activates the proteasome for RNF2 degradation.

**FIG 9.**
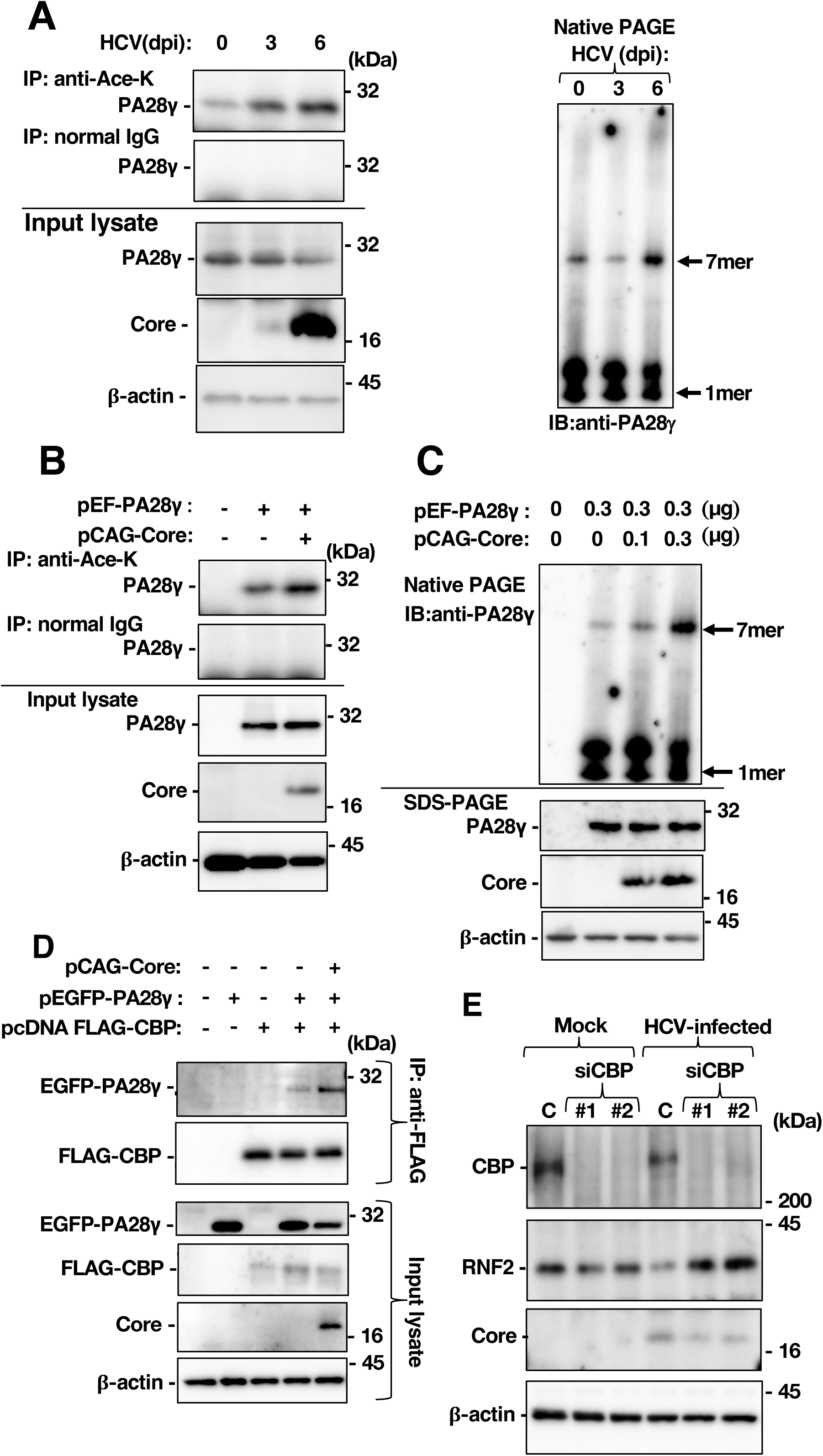
HCV core proteins induce the acetylation and activation of PA28γ via the KAT CBP, leading to a decrease in the RNF2 protein level. (A) Parental Huh7OK1 cells and PA28γ-knockout Huh7OK1 cells were infected with HCVcc at an m.o.i. of 0.5 and harvested at 0, 3 and 6 dpi. These cell lysates were subjected to immunoprecipitation using rabbit anti-Ace-K IgG or normal rabbit IgG. The resulting precipitates and cell lysates were subjected to Western blotting. (B) The cell lysates described in panel A were subjected to Western blotting to analyse the formation of PA28γ heptamers. (C) PA28γ-knockout Huh7OK1 cells were transfected with pEF PA28γ (0.3 µg/sample) and/or pCAG Core (0.3 µg/sample). Total DNA was supplemented with the empty plasmid to a final amount of 1.0 µg per sample. The cells were harvested at 48 h post-transfection. These cell lysates were subjected to immunoprecipitation using rabbit anti-Ace-K IgG or normal rabbit IgG. The resulting precipitates and cell lysates were subjected to Western blotting. (D) The cell lysates described in panel C were subjected to Western blotting to analyse the formation of PA28γ heptamers. (E) PA28γ-knockout Huh7OK1 cells were transfected with pEGFP PA28γ (0.3 µg/sample), pcDNA3.1 FLAG-CBP (0.3 µg/sample), and/or pCAG Core (0.3 µg/sample). Total DNA was supplemented with the empty plasmid to a final amount of 1.0 µg per sample. The cells were harvested at 48 h post-transfection. The resulting cell lysates were subjected to immunoprecipitation and Western blotting. (E) Huh7OK1 cells were transfected with control siRNA (c), siCBP#1 (#1) or siCBP#2 (#2). The transfected cells were infected at 16 h post-transfection with HCVcc at an m.o.i. of 0.5. The infected cells were harvested at 3 dpi. The resulting cell lysates were subjected to Western blotting.

## DISCUSSION

Alterations in gene expression patterns in HCV-infected cells or liver tissue of individuals with hepatitis C may be involved in the pathogenic mechanism of HCV (34). We previously reported that HCV infection impaired PRC1-dependent monoubiquitination of histone H2A at K119 on HOX alleles through degradation of RNF2, which subsequently induced the expression of HOX genes (10). The data in this previous report also suggested that RNF2, a catalytic component of PRC1, is degraded in response to HCV infection or core protein expression and that this degradation is dependent on the proteasome but not on polyubiquitination (10). However, the mechanism by which the core protein induces the degradation of RNF2 after infection has not been elucidated. Furthermore, we previously reported that PA28γ is responsible for the core protein-related pathological changes. The PA28 family members, namely, α, β and γ, form heptamers that bind to the 20S proteasome and then stimulate polyubiquitin-independent proteasome activity. The PA28γ homoheptamer activates the 20S CP for the degradation of PA28γ-binding proteins. In mice, HCV core protein expression induces the development of hepatic steatosis, HCC and type II diabetes mellitus (5, 9, 23), which are highly prevalent in hepatitis C patients (35), and these core protein-related pathological conditions are reversed by PA28γ knockout in mice (22, 23). The data presented in our previous study also suggest that PA28γ binds the core protein and then induces the degradation of the unfolded core protein to promote viral particle production (20, 21). However, the mechanism by which the core protein induces HCV-related pathological changes via a PA28γ-related pathway remains unknown. In this study, the results of the yeast two-hybrid screen using PA28γ as bait revealed that RNF1 and RNF2 were the candidate PA28γ-binding partners. PA28γ bound both RNF1 and RNF2 (Fig. 1) and accelerated RNF2 degradation but only weakly affected RNF1 degradation, leading to the upregulation of HOX gene expression (Figs. 2 to 5). Furthermore, PA28γ promoted the proteasome-dependent degradation of RNF2 to impair the monoubiquitination of histone H2A (Figs. 4 to 6). Liu et al reported that CBP-induced acetylation of PA28γ at K195 resulted in homoheptamerization of PA28γ and subsequently stimulated polyubiquitin-independent proteasome activity (32). The data shown in Figs. 8 and 9 support the idea that CBP-dependent acetylation of PA28γ leads to heptamerization of PA28γ (Fig. 8) and suggest that core protein expression promotes the acetylation of PA28γ via CBP activity in infected cells (Fig. 9). Taken together, these data suggest that after HCV infection, the core protein reinforces the interaction between CBP and PA28γ, induces PA28γ heptamerization, and subsequently stimulates polyubiquitin-independent proteasome activity for RNF2 protein degradation. The data presented in this study also suggest that PA28γ expression is required for the reduction in H2Aub in HCV-infected cells. Identifying the other loci and genomic regions at which PA28γ reduces the level of H2Aub is an important and challenging task necessary for understanding the physiological and pathological functions of PA28γ in hepatitis C pathogenesis. A model of the mechanism by which HCV infection facilitates the degradation of RNF2 via a PA28γ-dependent proteasome pathway to impair histone H2A monoubiquitination and induce HOX gene expression is shown in Fig. 10.

**FIG 10.**
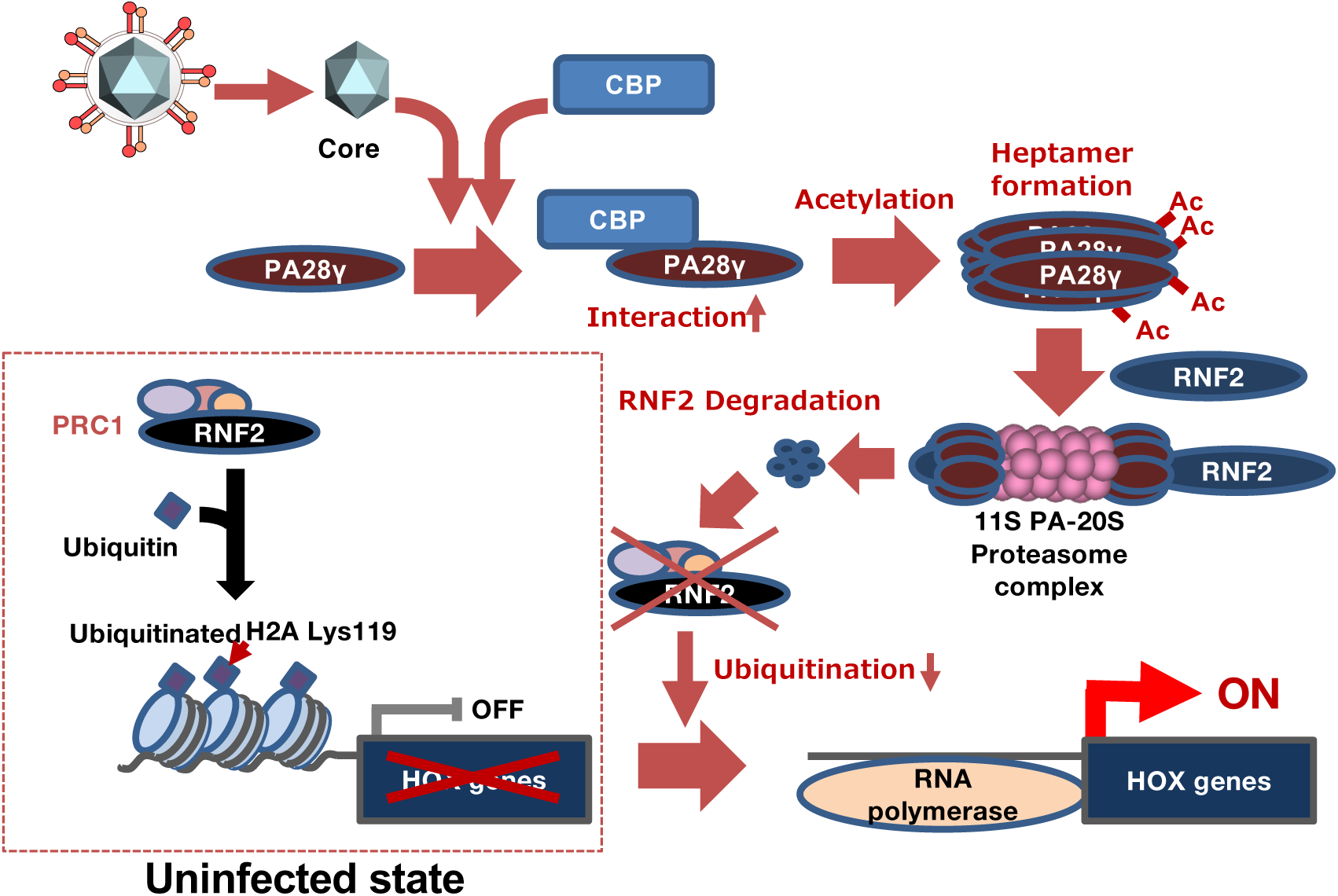
Schematic diagram of the mechanism by which HCV infection accelerates the activation of the PA28γ-20S CP complex for the degradation of RNF2, the reduction in H2Aub, and the induction of HOX genes. Most HOX alleles are silenced in uninfected hepatocytes (10). When hepatocytes are infected with HCV, the core protein reinforces the interaction between CBP and PA28γ to increase PA28γ acetylation and heptamerization. The PA28γ heptamer recruits RNF2 and activates the 20S CP to degrade RNF2. The degradation of RNF2 results in a decrease in PRC1 activity and impairs the monoubiquitination of histone H2A. The reduction in H2Aub results in the formation of open chromatin in the promoter regions of HOX alleles and the transcriptional reactivation of silenced HOX genes.

Several reports indicate that the core protein promotes the degradation of p21 (36, 37), whereas other reports suggest that the core protein induces G1 arrest after common bile duct ligation, alters the cell cycle in hepatocytes and is associated with increased p21 expression (38). PA28γ was reported to bind cyclin kinase inhibitors, such as p21 and p19, to promote their degradation via the PA28γ-20S CP complex and then to induce S phase entry (18, 39). These observations and the data presented in the current study suggest that the core protein may regulate the cell cycle via the control of the PA28γ-dependent proteasome. In addition, HOX protein expression induced by the core protein may promote the transcriptional activation of genes associated with degradation of cell cycle inhibitors. HOXA10 overexpression was reported to promote the expression of S-phase kinase-associated protein 2 (SKP2), which contributes to the degradation of the cell cycle inhibitor p27 (40). The upregulation of HOX gene expression in HCV-infected hepatocytes may be associated with the promotion of cell cycle progression via transcriptional and/or posttranslational regulation of cell cycle inhibitors.

The induction of HOX genes may contribute to carcinogenesis by regulating cell growth, survival, migration and invasion in addition to regulating the cell cycle. Several HOX genes have been reported to be dysregulated in a tissue-specific manner. HOXB13 is upregulated in breast cancer, whereas its expression is suppressed in prostate cancer (41, 42). HOXD10 is upregulated in breast, gastric, hepatocellular, colorectal, bladder and prostate cancers (43). HOXA9 is upregulated in acute lymphocytic leukaemia and epigenetically silenced in lung cancers (44, 45). HOXA5, HOXA7, HOXA13 and HOXD3 are highly expressed in HCC tissues compared to normal tissues (46). Notably, HOXA13 and HOTTIP were found to be highly expressed in the same neoplastic hepatocyte populations and in liver cancer cell lines (46, 47). HOTTIP, a long noncoding RNA (lncRNA) located contiguous to the HOXA13 gene, directly interacts with WDR5/MLL complexes to induce an open chromatin configuration by driving histone H3 Lys 4 trimethylation, leading to the induction of HOXA genes, including HOXA13 (48); moreover, HOXA13 can induce the transcription of HOTTIP as well as genes related to cell growth and cell invasion (47, 49). High expression of both HOTTIP and HOXA13 was correlated with metastasis formation and poor patient survival in HCC, suggesting that HOTTIP/HOXA13 expression may be a possible marker for predicting clinical outcomes in HCC patients (47). Although HCV infection cannot potently induce HOXA13 gene transcription in Huh7OK1 cells (10), the discrepancies in the data between our previous study and the abovementioned *in vivo* study may be due to the high expression of HOXA13/HOTTIP in the Huh7 cell line, regardless of the HCV infection status (47). Another lncRNA, HOTAIRM1, was also reported to induce the expression of HOXA1, which could promote the expression of Nanog (50). Some HOX genes associated with carcinogenesis may be constitutively expressed via a positive feedback loop in hepatocytes after DAA treatment.

In this study, HCV infection reduced the level of RNF2 and reactivated the expression of silenced HOX genes. However, some clinical reports suggest that the RNF2 and HOX proteins are highly expressed in cancer tissues (51). The data presented in this study may reflect the status of HCV-infected hepatocytes during the early stage and before the premalignant state of liver cancer but not during the malignant state of liver cancer. The expression of some HOX genes has been reported to be maintained in the malignant state of HCC regardless of the high expression of PRC1 components, including RNF2 and PCGF4 (BMI1), suggesting that the expression of some HOX genes may be regulated by factors other than PRC1, such as lncRNAs (e.g., HOTTIP, described above), regardless of the high RNF2 activity at the malignant stage of HCC. Increased expression of RNF2 in HCC tissue was reported to synergistically regulate crosstalk among the histone modifications H2Aub, H3K27me3 and H3Kme3 to suppress E-cadherin transcription (52). HCV infection increases the RNF2 mRNA level but reduces the RNF2 protein level in HCV-infected hepatocytes (10) and may subsequently irreversibly trigger RNF2 transcription, which may be maintained even after viral eradication at the malignant stage of HCC.

The possibility remains that PA28γ activation in HCV-infected cells may be associated with viral particle production in these cells. The data presented in our early work revealed that PA28γ knockdown reduced the production of HCV viral particles in the supernatant but did not affect viral RNA replication (21). In this study, compared with that in wild-type cells, the core protein level was much higher in either infected or core protein-expressing PA28γ-knockout cells (Fig. 6A, 7A), while PA28γ knockout reduced the infectious titre in the culture supernatant but did not affect viral RNA replication (Fig. 6C). Nonfunctional or unfolded core proteins were reported to interact with PA28γ for their proteasome-dependent degradation (20, 53). A functional core protein may be engulfed by a nonfunctional or an unfolded core protein in the absence of PA28γ to suppress viral particle production. Alternatively, a reduction in PRC1 activity may affect the expression of host genes related to membrane trafficking or viral assembly. Further study is required to clarify the mechanism by which depletion of PA28γ reduces the production of infectious HCV particles.

An increasing number of studies have suggested that the PA28γ proteasome is associated with several kinds of viral infections. Coxsackievirus B3 infection was found to promote PA28γ-mediated proteolysis of p53 (54), suggesting that p53 degradation increases viral replication by reducing the inhibitory influence of p53 on viral replication. On the other hand, PA28γ-dependent proteasome activity was associated with an inhibitory effect on viral replication. Ko et al reported that PA28γ acted as a corepressor of human T-lymphotropic virus 1 (HTLV-1) p30 and was required for the suppression of viral replication and maintenance of viral latency (55). Regarding hepatitis B virus (HBV) infection, PA28γ was reported to be associated with HBV pathogenesis. The HBV X protein (HBx) activates HBV promoters to increase HBV RNA transcription, increase the abundance of covalently closed circular DNA (cccDNA) and promote the degradation of the host protein Smc5/6, which can inhibit HBV replication (56, 57). HBx plays an important oncogenic role in the liver in hepatitis B patients (58). For example, HBx promotes the expression of PA28γ to degrade p16 to promote progression through the G1 and M phases (59). In addition, PA28γ stabilizes the HBx protein by competitively inhibiting the binding of HBx to Siah-1 (60). HBx was reported to be polyubiquitinated by Siah-1 for proteasome-dependent degradation (61), and PA28γ competitively inhibited the binding of Siah-1 to HBx, leading to an increase in HBx stability. Thus, PA28γ may be an attractive target for the treatment of hepatitis B and C as well as other viral diseases.

In summary, our data presented herein indicate that HCV infection or core protein expression reinforces the interaction between CBP and PA28γ and promotes PA28γ acetylation and heptamerization to activate the proteasome for RNF2 degradation and that a reduction in the RNF2 protein level impairs the monoubiquitination of histone H2A at K119 in the promoter regions of HOX alleles. The reactivation of silenced HOX genes may be associated with HCV-related hepatocellular carcinogenesis. PRC1-dependent histone H2A monoubiquitination silences non-HOX genes (62) and positively or negatively regulates metabolic and developmental gene expression in association with RNA polymerase (63). HCV infection may lead to regulatory effects on various genes via the PCR1-H2Aub pathway during acute and/or chronic infection. Thus, elucidating the mechanism by which HCV infection can regulate PRC1 activity to regulate genes other than HOX genes is important for clarifying the molecular mechanism of hepatitis C pathogenesis and for the development of therapies for hepatitis C.

## MATERIALS AND METHODS

### Cell lines, HCV strain

The HCV strain JFH-1, which belongs to genotype 2a (64), was used as the cell culture-adapted HCV (HCVcc) strain in this study. The viral RNA encoded in the plasmid pJFH1 was transcribed and introduced into Huh7 OK1 cells according to the method reported by Wakita et al. (64). The cell infection procedure was reported previously (10) and is described in detail in each figure legend. The Huh7 OK1 cell line, a subclone derived from Huh-7 cells that is highly permissive to infection with the JFH-1 strain, was cultured as reported previously (65). The human embryonic kidney 293T cell line was purchased from the American Type Culture Collection (Manassas, VA, USA). The lysine deacetylase inhibitor TSA and KAT inhibitor C646 were purchased from Sigma‒ Aldrich Japan (Tokyo, Japan).

### Plasmids

The gene encoding PA28γ was subcloned and inserted into the pGBKT7 vector (resulting in the pGBKT7PA28γ plasmid). N-terminal HA-tagged RNF1 or RNF2 was amplified by polymerase chain reaction (PCR) from human liver cDNA using *KOD* DNA polymerase (TOYOBO, Osaka, Japan) and was then inserted into pCAGGS-PM3. The DNA fragment encoding PA28γ was cloned and inserted into pEF pGKpuro and pEF FLAG Gs pGKpuro as reported previously, with the resulting plasmids designated pEF PA28γ and pEF FLAG-PA28γ, respectively (21), as well as into pEGFP-C3 (Clontech Takara, Tokyo, Japan). The plasmid encoding the HCV core protein (amino acids 1-192) was used for cloning and insertion of the corresponding DNA into pCAGGS-PM3 as described previously, and the resulting plasmid was designated pCAG-Core (21). N-terminal FLAG-tagged CBP/p300 cDNA was amplified by PCR and was then inserted into pcDNA3.1 (Invitrogen Thermo, Carlsbad, CA).

### Yeast two-hybrid screening

The bait plasmid pGBKT7PA28γ was introduced into *Saccharomyces cerevisiae* AH109 cells. Yeast cells containing pGBKT7PA28γ were grown in yeast extract-peptone-dextrose medium and transfected with a human foetal brain plasmid library based on pACT2 (Clontech Takara). Clones (22.4 × 10^6^) generated with the library were screened (Clontech Takara, Tokyo, Japan). Yeast clones containing pGBKT7-53 and pGADT7-T (Clontech) were used as positive controls, while yeast clones containing pGBKT7 and pGADT7 were used as negative controls. Yeast colonies grown on dropout plates lacking tryptophan, leucine, histidine, and adenine were inoculated on two fresh dropout plates lacking leucine and tryptophan. One of the two plates was used for a β-galactosidase assay according to the method of Duttweiller (66), and the other plate was stored at 4 °C as a master plate. One of the 556 dropout plate-positive clones exhibited dark blue staining on a β-galactosidase assay plate to the same extent as the positive control and was called C1-26. The plasmid in Cl-26 cells encoded the RNF1 gene. We amplified the genes encoding RNF1 and the RNF1 homologue RNF2 from the total human liver cDNA by PCR and then inserted the sequences into several plasmids as described above. We also isolated the yeast clone encoding PA28γ by this screening method, which supported the accuracy of this screening method.

### Transfection, immunoblotting, immunoprecipitation, and gene silencing

Plasmid DNA was transfected into Huh7OK1 or 293T cells by using TransIT^®^-LT1 (Mirus Bio, Madison, WI). Lysate preparation and immunoprecipitation were carried out as described previously (21). The cell lysates were subjected to 5-20% sodium dodecyl sulfate‒polyacrylamide gel electrophoresis (SDS‒PAGE) for standard protein analysis or to native PAGE using a NativePAGE^TM^ 3‒12% mini protein gel and a Sample Prep Kit (Thermo Fisher) for analysis of the heptamerization of PA28γ. The proteins in the gel were transferred onto polyvinylidene difluoride membranes (Merck Millipore, Billerica, MA). After protein transfer, the membranes were incubated first with an appropriate primary antibody and then with a horseradish peroxidase-conjugated rabbit or mouse IgG as a secondary antibody, immersed in Super Signal West Femto (Thermo Fisher Scientific, Rockford, IL) and visualized using an LAS 4000 Mini imaging system (Cytiva, Marlborough, MA). The small interfering RNAs (siRNAs) targeting RNF2 and the control siRNA (siControl Non-targeting siRNA #2, Dharmacon®) were purchased from Thermo Scientific (Brebières, France) and were introduced into cells by using Lipofectamine RNAiMax (Thermo Fisher Scientific). The Silencer® siRNAs with ID numbers s3495 and s3497 were purchased from Thermo Fisher Scientific and are designated CBP#1 and siCBP#2, respectively, in this study. The CHX chase assay was carried out by the method reported by Kao et al (67).

### Reverse transcription–quantitative polymerase chain reaction (RT‒qPCR) and reverse transcription–semiquantitative polymerase chain reaction (RT–sqPCR)

Total RNA and first-strand cDNA were prepared and then evaluated by RT‒qPCR as described previously (10). The expression levels of HCV RNA and of each host mRNA were normalized to that of GAPDH mRNA. HCV RNA and GAPDH mRNA were amplified using primer pairs as described previously (21, 65). Each PCR product was confirmed to be detected as a single band of the correct size by agarose gel electrophoresis. The amount of HCV in the culture supernatant was estimated as the copy number (10). HOX gene mRNA levels were measured by RT–sqPCR or RT‒ qPCR (68) using the appropriate primer pairs as described previously (10). Takara Emerald (2x;Takara Co.) was used for RT–sqPCR. Data acquisition was performed for 25, 30, 35, and 40 cycles to confirm that the plateau phase had not been reached.

### Evaluation of immunofluorescence staining in cells by microscopy

Huh7OK1 wild-type or transfected cells were infected with HCVcc, passaged twice every 4 days, seeded at 0.5 × 10^4^ cells per well on a glass cover slip and incubated at 37 °C for 24 h. The cells were washed twice with phosphate-buffered saline (PBS) and fixed with 4% paraformaldehyde at room temperature for 20 min. After fixation, the cells were washed twice with PBS, permeabilized by incubation for 15 min at room temperature in PBS containing 0.3% saponin, and then incubated in PBS containing 3% bovine serum albumin (PBS-BSA) to block nonspecific signals. These cells were stained with 50 µM 4’,6-diamidino-2-phenylindole (DAPI) and incubated at 4 °C overnight in PBS-BSA containing rabbit anti-RNF2 IgG (clone D22F2, Cell Signaling Technology, Denver, MA) and mouse anti-NS5A IgG (clone 9E10) or rabbit anti-H2Aub IgG (clone D27C4, Cell Signaling Technology) and mouse anti-HCV core protein IgG (clone B2, ANOGEN, Mississauga, Canada). The cells were washed three times with PBS-BSA and incubated at room temperature for 2 h in PBS-BSA containing appropriate Alexa Fluor (AF)488 or AF594-conjugated secondary antibodies (Thermo Fisher Scientific). The cells were washed three times with PBS-BSA and observed using a confocal laser scanning microscope (FV1000, Olympus, Tokyo) or fluorescence microscope (BZ-9000 and BZ-X800: Keyence, Osaka, Japan).

After reacting with each the first antibody, PA28γ and RNF2 were detected using AF488- and AF594-conjugated secondary antibodies, respectively. Nuclei were stained with DAPI. Localization signals of PA28γ and RNF2 were obtained as three-dimensional (Fig. 1C) and two-dimensional images (Figs. 1B and 4B) using all-in-one fluorescence microscope (BZ-X800, KEYENCE, Osaka, Japan) equipped with a Plan Apochromat 40x objective (NA0.95, BZ-PA40, KEYENCE, Osaka, Japan) and optical sectioning module (BZ-H4XF, KEYENCE, Osaka, Japan). Green fluorescence was detected using a GFP filter (ex:470/40nm, em:525/50nm, dichroic:495nm, OP-87763, KEYENCE, Osaka, Japan), while red fluorescence was detected using a Texas Red filter (ex:560/40nm, em:630/75nm, dichroic:585nm, OP-87765, KEYENCE, Osaka, Japan). Blue fluorescence was detected using a DAPI filter (ex:360/40nm, em:460/50nm, dichroic:400nm, OP-87762, KEYENCE, Osaka, Japan). Images were acquired as continuous images using the sectioning function. Fluorescence was measured for the image at the center depth of the Z-axis of the acquired serial images, and each signal intensity of the same coordinate pixel within the DAPI-positive region (nuclear region) was compared and evaluated using a fluorescence microscope and the corresponding software (BZ-H4: Keyence, Osaka, Japan). Co-localization of PA28γ and RNF2 (Fig. 4B) was determined using the function evaluating co-localization included in a confocal laser microscope FV1000.

### ChIP assay

ChIP assays were carried out according to the method reported by Sakurai et al (69) with modifications as described previously (10). Huh7OK1 cells were infected with HCVcc and passaged twice every 4 days. HCV-infected cells and mock-infected cells were harvested 8 dpi. DNA‒protein complexes were prepared as described previously (10). qPCR was carried out as described above without a reverse transcription step.

### Gene knockout with the CRISPR/Cas9 system

PA28γ-knockout Huh7OK1 cell lines were established according to the method reported by Fujihara et al. (70). A PA28γ-targeting sequence (5’-ATGGGATGCTGAAAAGCAAC-3’), which is located in the exon 5, was used for the construction of the pX330 plasmid encoding the guide RNA. The targeted DNA regions were amplified from the genomic DNA of Huh7OK1 cells with the primer pair 5’-ACTGAGGATCCATGTTTTAGACGCTCATCTGTAGTTC-3’/5’-GATATCGAATTCCAACCCAGGAGGCAGAGGTC-3’ and were subsequently inserted into pCAG EGxxFP. Both pX330 and pCAG EGxxFP were purchased from Addgene (Cat Nos. #42230 and #50716; Cambridge, MA). Huh7OK1 cells were transfected with these plasmids using Lipofectamine LTX (Thermo Fisher Scientific). GFP-positive cells were isolated using a FACSAria^TM^ cell sorter (BD Bioscience) 48 h post-transfection. Single-cell clones were established by a colony isolation technique. The deleted regions of each allele of the PA28γ gene were confirmed by direct sequencing. The PA28γ mRNA transcribed from each allele in the PA28γ-knockout cell line lacked the sequence 5’-GCTGAAAAGCAACCAGCAGC or ACCAGCAGCTG-3’, corresponding to the region on the downstream of amino acid residue 7 or 3. The expression of PA28γ was confirmed by immunoblot analysis and direct sequencing. There appears to be no difference between wild-type and knockout cells with respect to cell proliferation.

### In vitro proteasome assay

The plasmids encoding HA-RNF1 and HA-RNF2 were transfected into PA28γ-knockout Huh7OK1 cells (5 x 10^5^ cells) and then harvested 48 hr. The cell lysates were prepared in 1 ml of NP-40 Lysis buffer (50 mM Tris 7.4 pH, 150 mM NaCl, 1% NP-40, 10% glycerol, 1x protease inhibitor cocktail). HA-RNF1 or HA-RNF2 was then isolated by immunoprecipitation using an anti-HA antibody for *in vitro* proteasome assays. The in vitro proteasome assay was conducted using purified 20S proteasome (Enzo Life Science, Farmingdale, NY) and proteasome activator 11S γ subunit (Enzo Life Science). The immunoprecipitates containing HA-RNF1 or HA-RNF2 (one-twentieth of the total immunoloprecipitates) were mixed on ice with 20S CP (50 ng protein) with or without the 11S PA consisting of PA28γ (28 ng protein) in proteasome assay buffer (25 mM Tris-HCl pH7.4, 10% glycerol, 10 mM MgCl_2_, 1 mM DTT, 2 mM ATP). The mixture was incubated at 37 °C and stopped at 0, 10 or 30 min by boiling after adding SDS-sample buffer. The resulting mixtures were analysed by Western blotting. Intensity of RNF1 and RNF2 protein signals were measured using Image Gauge software (GE health care).

### Statistical analysis

The measured values are shown as the means ± standard deviations (SDs). Normality tests were performed by the Shapiro-Lorc test and Q-Q plots. Correlation tests were performed by Pearson’s correlation test for samples that showed a normal distribution, and by Spearman’s rank correlation test for the others. The statistical significance of differences in the means was determined by Student’s *t* test. A *p*-value less than 0.05 was considered statistically significant (*: *p* < 0.05, **: *p* < 0.01).

## Acknowledgements

We thank T. Wakita, F. V. Chisari and R. Bartenschlager for providing plasmids and cell lines. We also thank M. Mori for her secretarial work and C. Endoh for technical assistance. This work was supported by Grants-in-Aid from the Japan Agency for Medical Research and Development (24fk0210109), JST Moonshot R&D (JPMJMS2025), and scholarship donations from Teijin Co. Ltd and Yakult Co., Ltd.

